# Transcriptional control of adipogenesis by PATZ1

**DOI:** 10.1101/2022.05.24.493273

**Authors:** Sanil Patel, Njeri Sparman, Shani Sadeh, Jeixin Wang, Julian P. Whitelegge, Susan K. Fried, Hironori Waki, Claudio J. Villanueva, Marcus Seldin, Shinya Sakaguchi, Wilfried Ellmeier, Peter Tontonoz, Prashant Rajbhandari

## Abstract

White adipose tissues (WAT) play a central role in lipid storage and systemic energy, lipid, and glucose homeostasis. Understanding the intricacies of adipocyte formation could inform therapies for obesity and metabolic disorders. We have identified the POZ/BTB and AT Hook Containing Zinc Finger 1 (PATZ1) protein as an adipogenic transcription factor through an unbiased high-throughput cDNA screen for modulators of adipogenesis. PATZ1 is expressed by both human and mouse adipocyte precursor cells (APC) and adipocytes, and its expression negatively correlates with obesity. In cell models, PATZ1 expression promotes adipogenesis through a mechanism dependent on protein-protein interaction and DNA binding. Adipose-specific ablation of PATZ1 in mice leads to decreased lipid storage and fat mass and protection from diet-induced obesity. Genome-wide PATZ1 DNA binding analyses using ChIP-Seq suggest that PATZ1 facilitates adipogenesis through interactions with transcription factor machinery at the promoter regions of key adipogenic factors and histone modifiers. Global PATZ1 protein interaction studies using immunoprecipitation-mass spectrometry (IP-MS) suggest that General Transcription Factor 2I (GTF2I) binding to PATZ1 forms a repressive complex, and knockdown of GTF2I augments PATZ1 adipogenic function. Collectively these findings identify PATZ1 as a regulator of the adipocyte differentiation programs and advance our understanding of the complex transcriptional mechanisms underlying adipose tissue development and homeostasis.

## Main

Adipocytes, along with osteoblasts, myocytes, and chondrocytes are derived from pluripotent mesenchymal stem cells (MSCs) ^1^. The formation of adipocytes from precursor MSCs involves a complex and highly orchestrated program of gene transcription ^2^. Adipogenesis can be divided into two phases: commitment and terminal differentiation. The commitment phase involves dynamic integration of cytoarchitecture, signaling pathways, and transcriptional regulators. Multiple signals from bone-morphogenetic factor (BMP), transforming growth factor (TGF), fibroblast growth factor (FGF), and WNT influence stem cell conversion to adipocytes ^3^. Zinc-finger protein 243 (ZFP423), TCF7-like 1 (TCF7L1), and Krüppel-like factors (KLFs) are among the transcriptional regulators that have been shown to play essential roles in the determination of MSCs to form adipocyte precursors ^4-6^. After the commitment phase, a cascade of transcriptional events is activated that induces the expression of metabolic genes and adipokines associated with terminal adipocyte differentiation ^2^. Transcriptional events in the terminal differentiation phase have been well studied, and more than twenty different transcription factors have been identified as playing key roles. Among them, peroxisome proliferator-activated receptor gamma (PPAR*γ*) and CCAAT/enhancer-binding proteins (C/EBPs) are the primary drivers of gene induction during terminal differentiation ^2,7^.

PATZ1 has an N-terminal POZ/BTB domain for protein interaction, a central DNA binding AT hook domain, and a second, C-terminal, zinc finger (ZF) DNA binding domain ^8^. PATZ1 is an architectural transcriptional regulator that negatively or positively modulates the expression of genes by binding to promoter GC-rich regions. PATZ1 is known to regulate stem cell pluripotency and reprogramming, spermatogenesis, T-cell development, and cancer progression ^8-10^. PATZ1 interacts with other transcriptional regulators known to affect adipose homeostasis, including BCL6 ^11^, p53 ^10^, NCoR ^12^, Bach1 ^13 14-16^. However, the role of PATZ1 in metabolism and adipose biology is still unexplored. Here we present PATZ1 as a novel adipogenic transcription factor that promotes differentiation of adipocytes by interaction with the transcriptional machinery at the promoter regions of other adipogenic factors.

## Results

### A high-throughput screen of cDNA modulators of adipogenesis identifies PATZ1 as an adipogenic factor

To identify novel factors involved in adipocyte differentiation, we analyzed data from a cDNA library screen in which 10T1/2 cells were retro-transfected simultaneously with a luciferase reporter driven by the −5.4 kb *Fabp4* promoter and a collection of 18,292 individually spotted mammalian cDNA expression vectors (**Fig. 1A)** ^17^. The day after transfection, cells were treated with insulin and a PPARγ agonist (rosiglitazone) to induce adipogenic differentiation then luciferase activity was evaluated 4 days later. A number of cDNAs were identified as activators of aP2-driven luciferase activity. PPARγ emerged as the most potent activator, and several additional known adipogenic factors were also represented, including CCAAT/enhancer-binding proteins (C/EBPα and C/EBPδ), early B cell factor 1 (EBF1), and mitogen-activated protein kinase kinase 6 (MAPKK6). We recently reported adipogenic roles of paraspeckle protein 1 (PSPC1) and transducing-like enhancer protein 3 (TLE3), two other top candidates from this screen ^17-19^. To identify other cDNA modulators of adipogenesis, we tested three potentially novel regulators, PATZ1, nuclear transcription factor X-Box binding like 1 (NFXL1), and nuclear receptor coactivator 5 (NCOA5). The cDNAs encoding these genes were cloned into a retrovirus vector and stably overexpressed in 10T1/2 cells, which were then differentiated into white adipocytes. Among others, cells stably expressing PATZ1 showed highest adipogenic potential as indicated by Oil-Red-O staining (**Fig.1A**).

**Figure 1.**
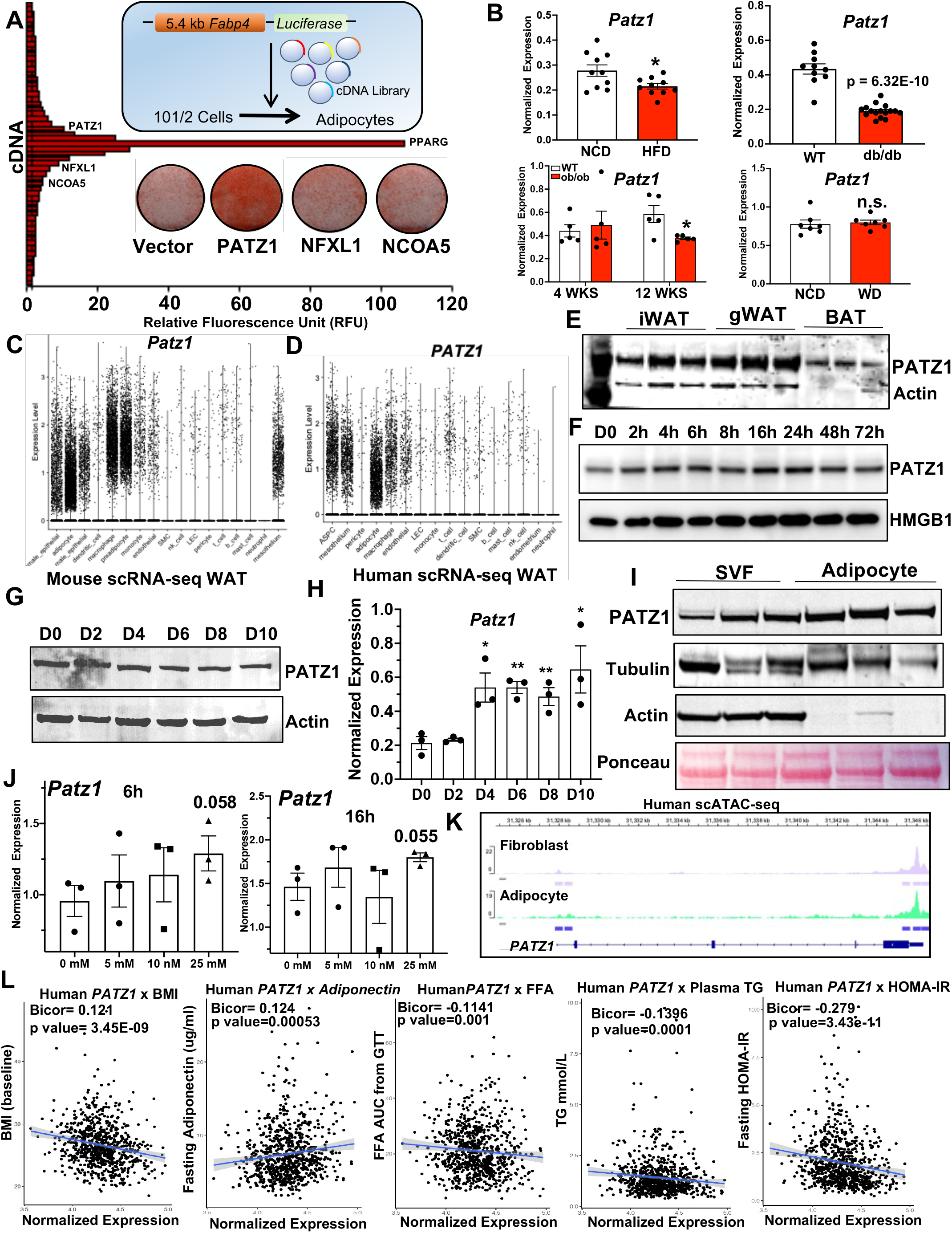
Identification of PATZ1 as an adipogenic factor. **A)** Diagram of cDNA library screen, plot of relative fluorescence Unit (RFU), and top hits of cDNA including PATZ1, NFXL1, and NCOA5. Oil-red-O staining of D6 differentiated 10T1/2 cells stably expressing Vector, PATZ1, NFXL1, and NCOA5. **B)** Real-time PCR analysis of PATZ1 mRNA in mice fed normal chow diet (NCD), high fat diet (HFD) for 12 weeks, or 10-week-old *db/db*, 4 week- or 12 week-old *ob/ob* mice, or mice fed western diet (WD) for 10 weeks. **C and D)** Violin plot of scRNA-seq ^20^ cluster analysis showing levels of PATZ1 mRNA in different cell types from mouse WAT (C) and human WAT (D). **E**) Western blot showing PATZ1 protein levels in mouse iWAT, gWAT, and BAT. Actin was used as a loading control. **F**) Western blot showing PATZ1 levels in white-differentiated 10T1/2 cells for indicated times. HMGB1 was used as loading control. **G and H**) Western blot (G) and real-time qPCR (H) showing PATZ1 levels in white differentiated 10T1/2 cells for indicated times without rosiglitazone. Actin was used as loading control for G. **I)** Western blot showing PATZ1 levels in SVF and adipocyte fraction from iWATs of 10-week-old chow fed mice. Tubulin, Actin, and ponceau stain were used as loading controls. **J)** Real-time PCR analysis of PATZ1 mRNA in undifferentiated 10T1/2 cells treated with indicated concentrations of glucose for 6h and 16h. **K)** scATAC-seq^23^ peaks in human WAT adipocytes and fibroblast at the promoter region of human PATZ1 gene. **L)** Correlation of human WAT PATZ1 levels with indicated metabolic traits from METSIM study. *, p < 0.05; **, p < 0.01. Figures with bar graphs and western blot are from experiments with at least three independent repeats.

To understand how obesity impacts expression of PATZ1, we analyzed *Patz1* expression in normal chow diet (NCD), *ob/ob, db/db*, high-fat diet (HFD), and western diet (WD) fed mice. Consistent with the expression of adipogenic factors like PPARγ and adiponectin ^19^, we noticed decreased *Patz1* transcript levels in these models (**Fig. 1B)**. Single cell RNA-seq (scRNA-seq) data from mouse and human WAT showed that *Patz1* was expressed by both APCs and adipocytes (**Fig. 1C and 1D)** ^20^. PATZ1 protein was also expressed in both inguinal WAT (iWAT) and gonadal WAT (gWAT) and to a lesser extent in BAT (**Fig. 1E)**. Further, PATZ1 levels remained relatively unchanged during differentiation of 10T1/2 cells, with a slight increase at 24 h (**Fig. 1F**). Adipocyte differentiation in the absence of PPARγ ligand did not alter PATZ1 protein levels despite increased mRNA expression (**Fig. 1G and 1H)**. Consistent with the scRNA-seq data, fractionation of WAT showed that PATZ1 was expressed in both the SVF and adipocyte portions of WAT (**Fig. 1I**).

PATZ1 was reported to be upregulated during hyperglycemia in endothelial cells ^21^; accordingly, treating glucose-starved 10T1/2 cells with increasing concentrations of glucose showed an increase in *Patz1* levels (**Fig. 1J**). A recent study using transcriptomics analysis to identify transcription factors in the upstream gene regulatory networks that drive adipocyte formation in human ASCs identified PATZ1, among other transcription factors ^22^. Single cell assay for transposase accessible chromatin-seq (scATAC-seq) in human adipocyte showed an open chromatin configuration, consistent with active transcription of *PATZ1* in human adipocytes ^23^ (**Fig. 1K**). Prior studies of genome wide DNA methylation and mRNA expression patterns in human adipose tissue identified PATZ1 among genes for which both DNA methylation and expression correlated with BMI ^23^. We next determined the correlation of WAT PATZ1 levels with human metabolic traits within the METSIM ^24,25^ population data and found negative correlations of PATZ1 with body mass index (BMI), plasma free fatty acid (FFA) levels, plasma triglyceride (TG) levels, and insulin resistance (HOMA-IR). We found a positive correlation between PATZ1 and adiponectin levels (**Fig. 1L**). Altogether, the expression pattern of PATZ1 is consistent with a potential function in adipocyte gene expression.

### PATZ1 promotes white adipogenesis in vitro

To elucidate the role of PATZ1 in adipogenesis, we generated 10T1/2 cells stably overexpressing PATZ1 (PATZ1) or vector-expressing control cells (Vector) and differentiated them into white adipocytes for 8 days. We then performed global RNA-seq on the three independent replicates of differentiated Vector and PATZ1 cells. Principle component analysis (PCA) showed a similar clustering pattern of the replicates from both PATZ1 and Vector cells (**Fig. 2A**). The scatter plot of the total transcript analysis showed comparable expression patterns (R=0.989) between Vector_1 and Vector_2, and between PATZ1_1 and PATZ1_2 (**Fig. 2B**). Both groups of samples (Vector and PATZ1) had comparable overall distribution and density of the transformed data, implying that PATZ1 overexpression did not cause overt genomic transcript shifts during adipocyte differentiation (**Fig. 2C**). To probe for transcriptomic differences between Vector and PATZ1 cells, we performed differential gene expression (DEG) analysis and found 1,131 genes upregulated and 543 genes downregulated in PATZ1 cells compared to Vector cells (**Fig. 2D**). Volcano plot and heatmap analysis of DEGs between PATZ1 and Vector cells showed an increase in genes such as *Ephx2, Cfd, Plin1, Aqp7, Retn, Fabp4, and Adipoq* in PATZ1 cells (**Fig. 2E**). We isolated upregulated, downregulated, and similarly expressed DEGs and represented them as heatmaps and found a marked increase in “fat cell differentiation” pathway genes in PATZ1 cells (**Fig. 2F**). We also noticed an upregulation of “Brown fat cell differentiation” and “positive regulation of cold-induced thermogenesis” pathways in PATZ1 cells (**Fig. 2F**). Sub-setting global DEGs from the heatmap showed a high enrichment of adipogenic genes in PATZ1 cells (**Fig. 2G**). Gene ontology and pathway analysis from most RNA-seq are unbiased but could be driven by a few genes that may over-represent a specific pathway. Based on our subset of genes, we performed a rigorous test by binning DEGs in four clusters (A-D) and performing pathways analysis using individual genes to define clusters. The results were represented as a heatmap (**Fig. 2H**). Single gene cluster analyses were plotted as a tSNE plot, which showed a highly specific clustering of pathways (A-D) based on binning of highly expressed genes in PATZ1 cells (**Fig. 2I**). Cluster A pathways were upregulated in PATZ1 cells and showed a highly specific increase in the white fat cell differentiation program (**Fig. 2I bottom**).

**Figure 2.**
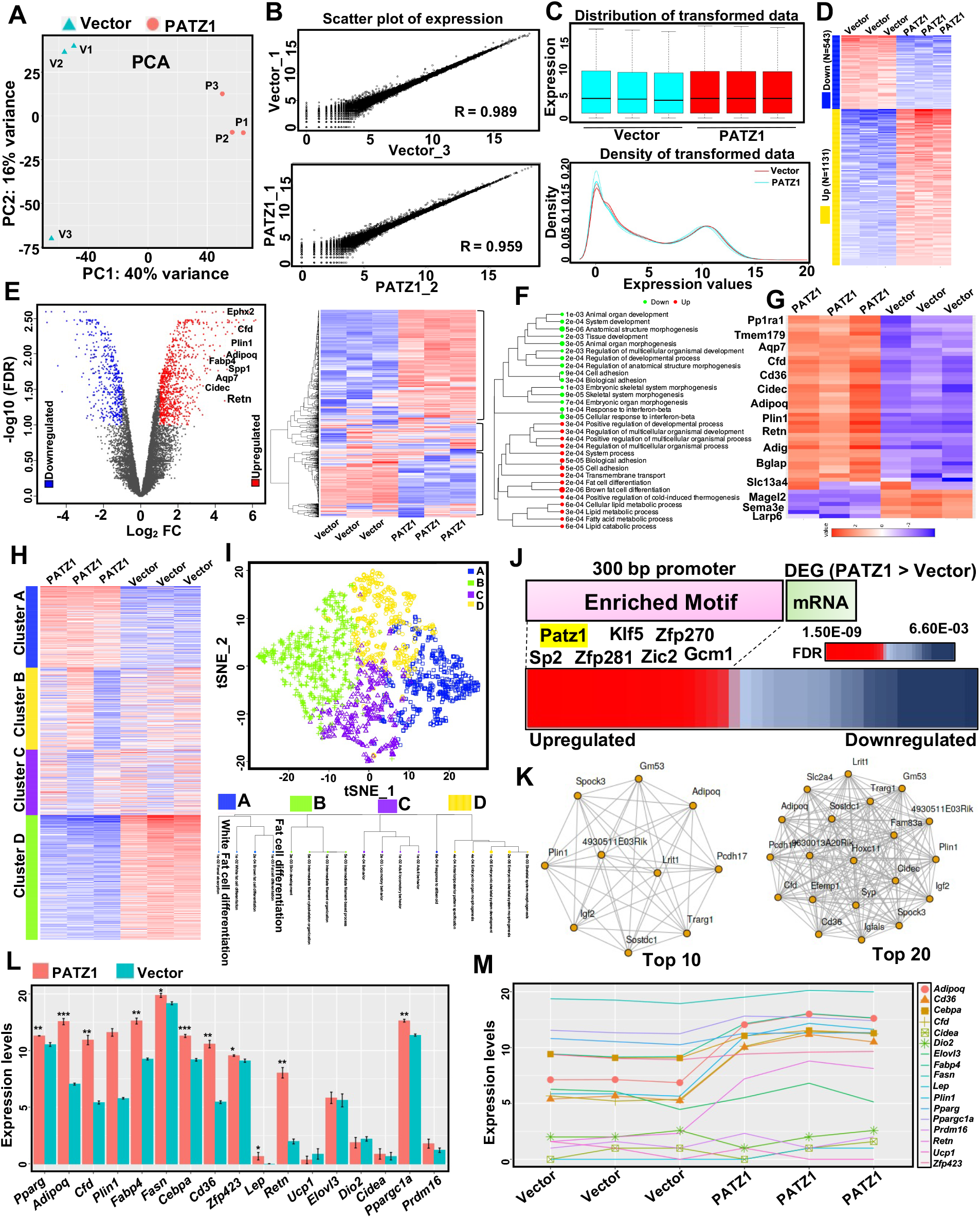
RNA-seq shows a highly specialized role of PATZ1 in adipogenesis. **A)** PCA of individual libraries generated from RNA isolated from vector (Vector) or PATZ1 overexpressing 10T1/2 differentiated for 8 days. **B)** Scatter plot of total gene expression correlation analysis of indicated libraries. **C)** Distribution and density of transformed expression data from indicated libraries from Vector and PATZ1 cells. **D)** Heatmap showing total genes upregulated and downregulated in Vector and PATZ1 cells. **E**) (Left) Volcano plot of gene expression differences between PATZ1 and Vector cells. The log2FC of PATZ1/Vector was plotted as a function of -log10 (FDR) with selected genes indicated as upregulated in PATZ1 cells. (Right) Heatmap showing upregulated, downregulated and similarly expressed genes in Vector and PATZ1 cells. **F)** GO analysis of top genes upregulated and downregulated in PATZ1 cells. **G)** Heatmap showing top binned genes that are upregulated and downregulated in PATZ1 and Vector cells. **H**) GO analysis represented as a heatmap and tSNE plot of the binned genes divided into four clusters (A-D) representing differentially enriched pathways in PATZ1 and Vector cells. **J)** TF motif analysis 300 bp upstream of DEGs that are greater in PATZ1 cells. **K)** Network analysis of top 10 and top 20 genes that are enriched in PATZ1 cells. L and M) Expression values of indicated genes represented as average (L) or per library (M) from PATZ1 or Vector cells. * p<0.05, **p<0.01, *** p<0.001, n.s. not significant.

We next calculated enriched transcription factor binding motifs at the promoter regions of DEGs upregulated in PATZ1 cells. We set 300 bp upstream of DEGs as promoter regions and found an enrichment of PATZ1 motifs (FDR = 3.4E-09) among other transcription factors (**Fig. 2J**). These data suggest a PATZ1 driven transcriptional program in PATZ1 cells. Network analysis showed an interaction between transcriptional programs of adipogenic genes when we binned both Top 10 and Top 20 DEGs (**Fig. 2K**). Finally, we performed a directed gene expression (Fragments Per Kilobase of transcript per Million mapped reads, FPKM) analysis of WAT genes and BAT genes in PATZ1 and Vector cells and found an increase in white adipocyte genes (**Fig. 2L**). Brown adipocyte genes were comparable between PATZ1 and Vector cells except for *Ppargc1a* in PATZ1 cells, which could be one of the drivers of the “brown fat differentiation” GO pathway annotation in **Fig. 2F**.

To further test the role of PATZ1 in adipogenesis, we overexpressed PATZ1 in another adipogenic cell line (3T3-L1) via retroviral transduction. Control and PATZ1-overexpressing 3T3-L1 cells were differentiated for 8 days using an adipogenic cocktail (See Methods). Oil-Red-O staining on D8 showed that PATZ1 expression increased the capacity of 3T3-L1 cells to differentiate (**Fig. 3A**). Real-time quantitative PCR (qPCR) during the differentiation time course (D0, D4, D6, D8) confirmed an increase in adipocyte genes in cells overexpressing PATZ1. We next tested if PATZ1 also promoted adipogenesis of primary APCs isolated from mouse and human WAT. We isolated primary SVF from mouse iWAT and treated confluent cells with either Control or PATZ1 retrovirus and differentiated the cells from D0-D11. As shown in **Fig. 3B**, PATZ1 overexpression also enhanced primary APC adipogenesis as judged by a marked increase in Oil-Red-O staining. Since PATZ1 is also expressed in human WAT adipocytes, we reasoned that PATZ1 may also be involved in human adipocyte derived stem cells (hADSC) differentiation. We used primary hADSC isolated from abdominal WAT of human subjects and performed adipogenic differentiation studies. We found that PATZ1 increased lipid-laden adipocytes and increased the expression of adipogenic genes such as *Pparg, Adipoq*, and *Plin1*.

**Figure 3.**
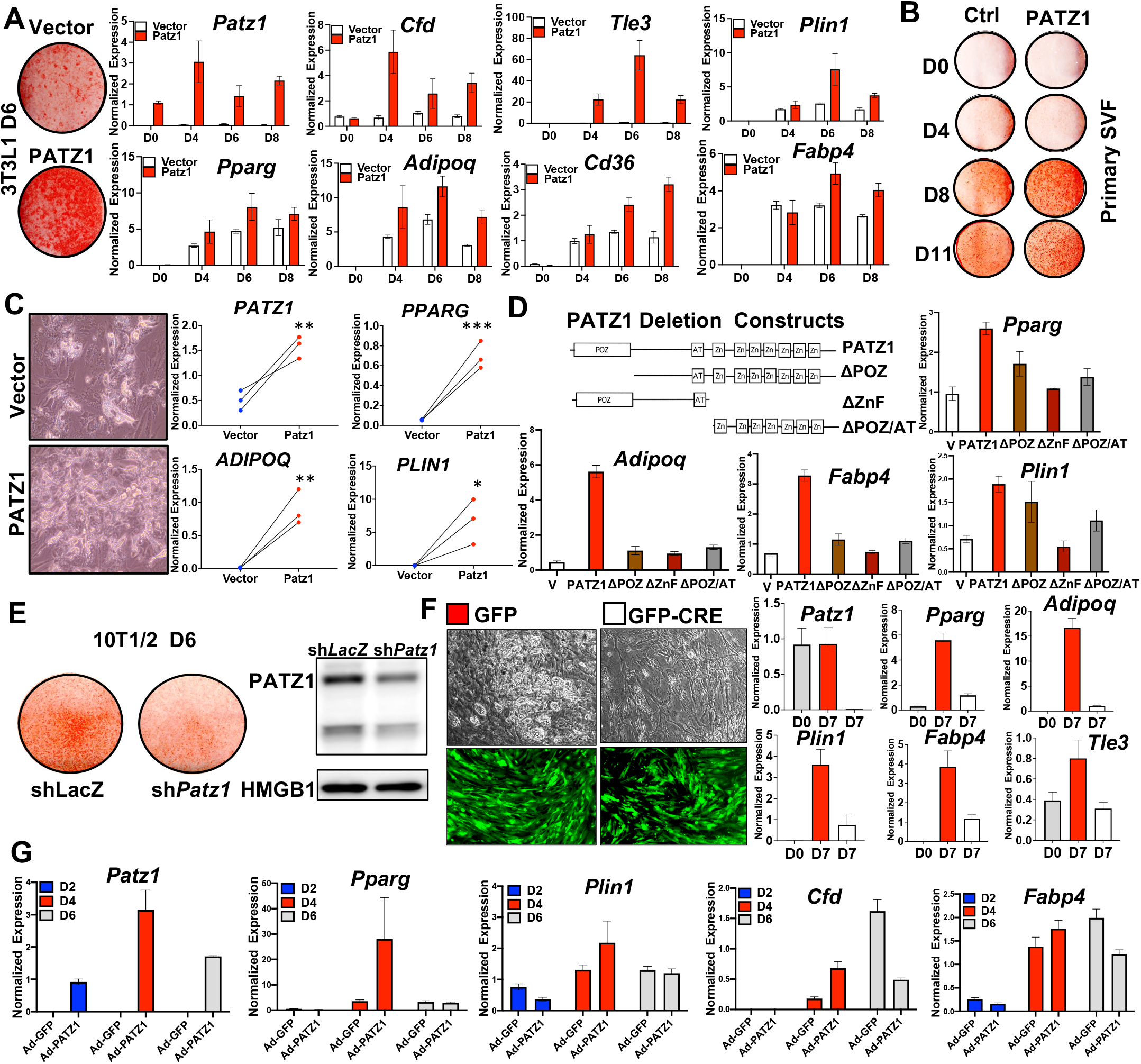
PATZ1 promotes adipocyte differentiation *in vitro*. **A)** Oil-Red-O staining and real-time qPCR analysis of control and PATZ1-overexpressing 3T3L1 cells differentiated for indicated days. **B)** Oil-Red-O staining of primary iWAT SVFs transduced with control or PATZ1 retrovirus and differentiated for indicated days. **C**) Bright field microscopy and real-time qPCR analysis of control and PATZ1 overexpressing primary human abdominal subcutaneous hADSCs differentiated for 12 days. **D)** Domain structure of PATZ1 constructs with indicated deletion and real-time qPCR showing indicated mRNA levels in D4 differentiated 3T3L1 cells stably expressing indicated PATZ1 constructs. **E)** Oil-Red-O lipid staining (right), and immunoblot for PATZ1 and HMG1 (left) in day 5 differentiated 10T1/2 cells stably expressing control LacZ shRNA (shLacZ) or PATZ1 shRNA (shPatz1). **F)** (Right) Bright field (top) and fluorescence at 509 nm (bottom) microscopy showing lipid droplets (red arrow; top) and GFP fluorescence (bottom) in 7 days differentiated SVFs derived from PATZ1F/F mice infected with adenovirus (Adeno) expressing GFP or GFP-Cre and (Left) qPCR showing mRNA levels of indicated genes from PATZ1 F/F SVFs at D0 or D7 infected with Adeno GFP or Adeno GFP-Cre. GFP levels (bottom panel) show comparable viral infection. **G)** Realtime qPCR of indicated genes from differentiated 10T1/2 hCAR cells transducted with GFP (Ad-GFP) or PATZ1 (Ad-PATZ1) adenovirus on indicated days. All experiments here are representative of at least there independent repeats. * p<0.05, **p<0.01, *** p<0.001, n.s. not significant.

PATZ1 is a modular protein consisting of a protein interaction POZ domain and two DNA binding AT hook and Zinc finger domains. To test the importance of these domains in PATZ1 function, we generated PATZ1 mutants lacking POZ domain (ΔPOZ), zinc finger domains (ΔZnF), or POZ and AT hook domains (ΔPOZ/AT) and generated 10T1/2 cells stably expressing these mutants. Our qPCR analysis on differentiated 10T1/2 cells showed that overexpression of full-length PATZ1 increased expression of *Pparg, Adipoq, Fabp4*, and *Plin1*. However, cells expressing all three mutants had markedly reduced expression of *Pparg, Adipoq, Fabp4*, and *Plin1*, suggesting that both protein interaction and DNA-binding functions of PATZ1 are important for its adipogenic function (**Fig. 3D**).

To test if PATZ1 loss affects adipogenesis, we generated 10T1/2 cells stably expressing control or *Patz1* shRNA. Results in **Fig. 3E** showed that PATZ1 loss reduced adipogenic potential of 10T1/2 cells via Oil-Red-O staining. We also isolated SVFs from iWAT from *Patz1f/f* mice and treated cells with GFP- or GFP-Cre expressing adenovirus. As shown in **Fig. 3F**, Cre-expressing cells had almost no visible lipid droplets and qPCR analysis showed a complete knockout of PATZ1 and decrease in adipocyte genes. To test the impact of PATZ1 exogenous overexpression during adipocyte differentiation, we infected adenoviral receptor-expressing (hCAR) 10T1/2 cell with GFP (Ad-GFP) or PATZ1 (Ad-PATZ1) adenovirus on D0, D2, D4, and D6 of differentiation. As shown in **Fig. 3G**, compared to stable overexpression models, PATZ1 transduction during differentiation did not cause noticeable changes in adipocyte genes, indicating a possible role of PATZ1 in priming cells during early stages of adipocyte differentiation. Our data show that PATZ1 enhances adipocyte differentiation, and both protein interaction and DNA binding functions are important.

### PATZ1 ablation protects mice from diet-induced obesity and adiposity

To test the *in vivo* role of PATZ1, we generated an adipose-specific knockout of PATZ1 (AdPATZ1KO) by crossing *Patz1*^*f/f*^ (PATZ1F/F) mice and *Adipoq*-Cre mice (**Fig. 4A**). On HFD, female AdPATZ1KO (fAdPATZIKO) weighed less than controls and were visibly leaner (**Fig. 4B and 4C)**. Upon visual inspection after laparotomy, the size of iWATs and gWATs were reduced compared to controls, with no visible changes in quadriceps (quad), BAT, liver, spleen, or kidney (**Fig. 4C)**. Body composition analysis using magnetic resonance imaging (MRI) showed reduced fat mass in fAdPATZ1KO mice with no change in lean mass (**Fig. 4D and 4E)**. Female mice also showed reduced iWAT and gWAT weights (**Fig. 4F**). To test whether PATZ1 adipogenic action was sex-dependent, we performed HFD studies and found that male AdPATZ1KO (mAdPATZ1KO) mice displayed reduced fat mass and WAT weights, and small, but statistically significant decreases in body weight compared to controls (**Fig. 4G-4J**). Histological analysis of WAT from AdPATZ1KO mice showed exceedingly larger adipocyte size compared to controls, indicating a potential increased lipid loading capacity of adipocytes to compensate for reduced adipogenesis. These data show that PATZ1 regulates *in vivo* adiposity and protects mice from diet-induced obesity.

**Figure 4.**
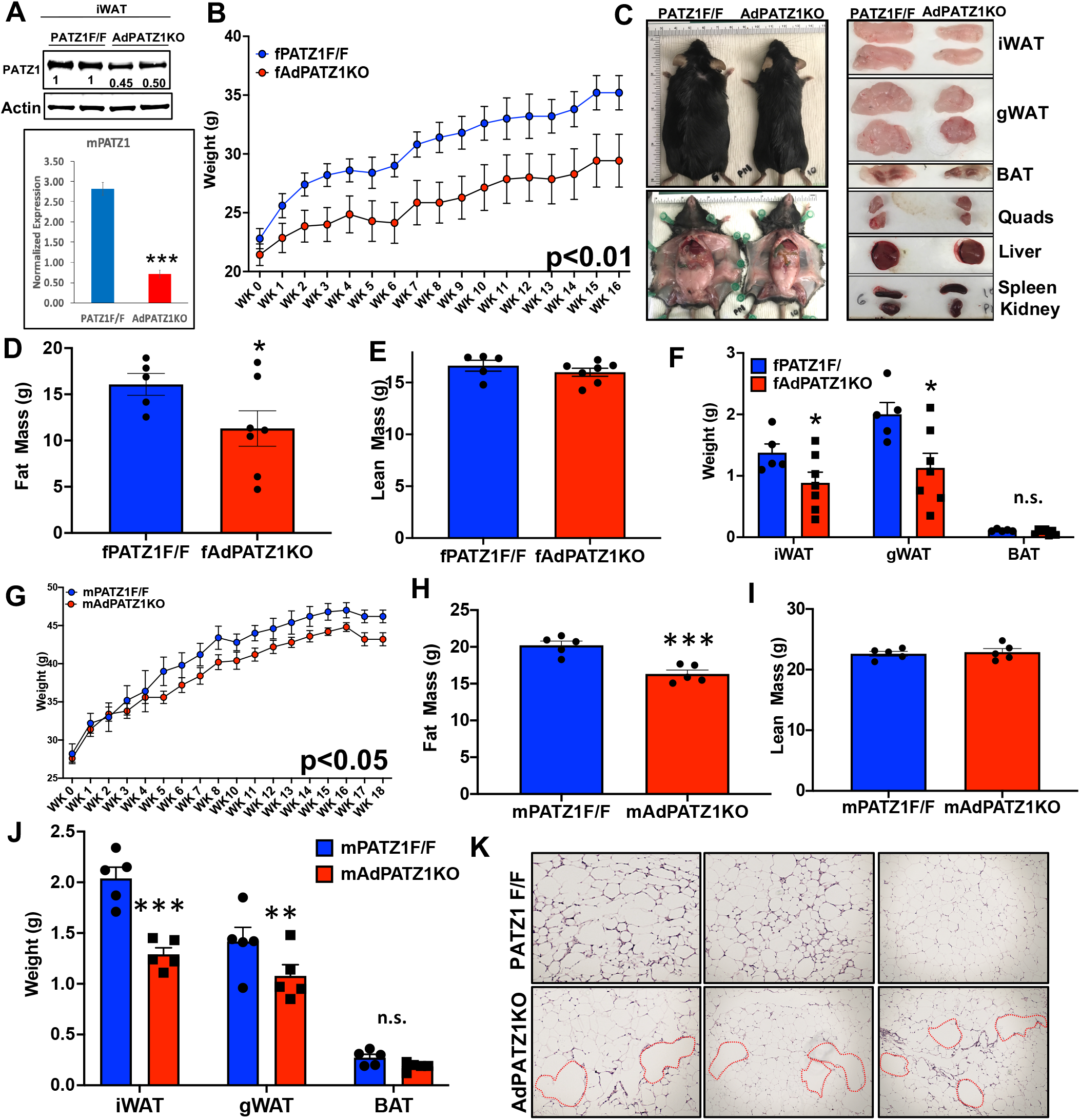
PATZ1 ablation protects mice from diet-induced obesity. **A)** Immunoblot and qPCR for PATZ1 in the iWATs of PATZ1F/F or AdPATZ1KO mice. Quantification of the bands showed ∼50% decrease in PATZ1 protein level in AdPATZ1KO mice. **B)** Body weight of AdPATZ1KO and PATZ1F/F female mice fed HFD for indicated weeks. **C)** (Left) External appearance of representative PATZ1F/F and AdPATZ1KO mice shown in B after 16 weeks on HFD (top) and representative PATZ1F/F and AdPATZ1KO mice after 16 weeks on HFD showing grossly reduced adipose tissue content in the absence of PATZ1 (bottom). (Right) Gross appearance of tissues from PATZ1F/F and AdPATZ1KO mice after 16 weeks on HFD. **D and E)** Body fat (D) and lean mass (E) in PATZ1F/F and AdPATZ1KO mice after 16 weeks on HFD determined by EchoMRI. **F)** Average weight of individual white and brown adipose fat pads from PATZ1F/F and AdPATZ1KO mice after 16 weeks on HFD. **G)** Body weight of AdPATZ1KO and PATZ1F/F male mice fed HFD for indicated weeks. **H and I)** Body fat (H) and lean mass (I) in PATZ1F/F and AdPATZ1KO mice after 18 weeks on HFD determined by EchoMRI. **J)** Average weight of individual white and brown adipose fat pads from PATZ1F/F and AdPATZ1KO mice after 18 weeks on HFD. **K)** Histology of gWAT from PATZ1F/F and AdPATZ1KO mice after 18 weeks of HFD. Red borders indicate large adipocytes in AdPATZ1KO mice. For all the figure panels, n = 5-7 per group, and represent cohorts from at least three independent studies. * p<0.05, **p<0.01, *** p<0.001, n.s. not significant.

### PATZ1 is dispensable for BAT development and function

BAT is a specialized adipose tissue that dissipates energy through non-shivering thermogenesis and controls systemic energy balance. We next asked whether PATZ1 was an adipogenic factor in BAT as well as WAT. We first generated stable BAT preadipocytes (PREBAT) overexpressing PATZ1 or Vector and differentiated these cells into mature brown adipocytes using a brown adipogenic cocktail (See methods). As shown in **Fig. 5A**, PATZ1 did not alter brown adipogenic potential of PREBAT cells, as shown by Oil-Red-O staining. However, transcript analysis showed upregulation of thermogenic genes such as *Ucp1* and *Cidea* without changes in adipocyte genes such as *Adipoq* and *Plin1* (**Fig. 5B**). These data suggested that PATZ1 could be involved in BAT function. To test the *in vivo* role of PATZ1 in BAT development and thermogenesis, we crossed PATZ1*f/f* mice with myogenic factor 5-Cre (*myf5-*Cre) to generate PATZ1 BAT knockdown (BAdPATZ1KO) in BAT precursor cells. We asked whether BAT *Patz1* deletion might result in altered systemic energy balance. Indirect calorimetry using metabolic chambers revealed no differences in oxygen consumption (VO2) and energy expenditure (EE) between BAdPATZ1KO and control mice at 10 weeks of age (**Fig. 5D**). BAdPATZ1KO mice showed a small reduction in lean mass and body weight compared to controls (**Fig. 5C**). Since MYF5 is also expressed in myogenic precursors, PATZ1 could potentially be involved in muscle development at an early age, but body composition analysis showed no differences between BAdPATZIKO and control mice at 24 weeks of age. (**Fig. 5E**). To test if BAdPATZ1KO mice showed age-dependent changes in systemic energy balance, we performed indirect calorimetry in 24-week-old BAdPATZ1KO and control mice. Metabolic chamber analysis at RT or cold showed no difference between the genotypes (**Fig. 5F**).

**Figure 5.**
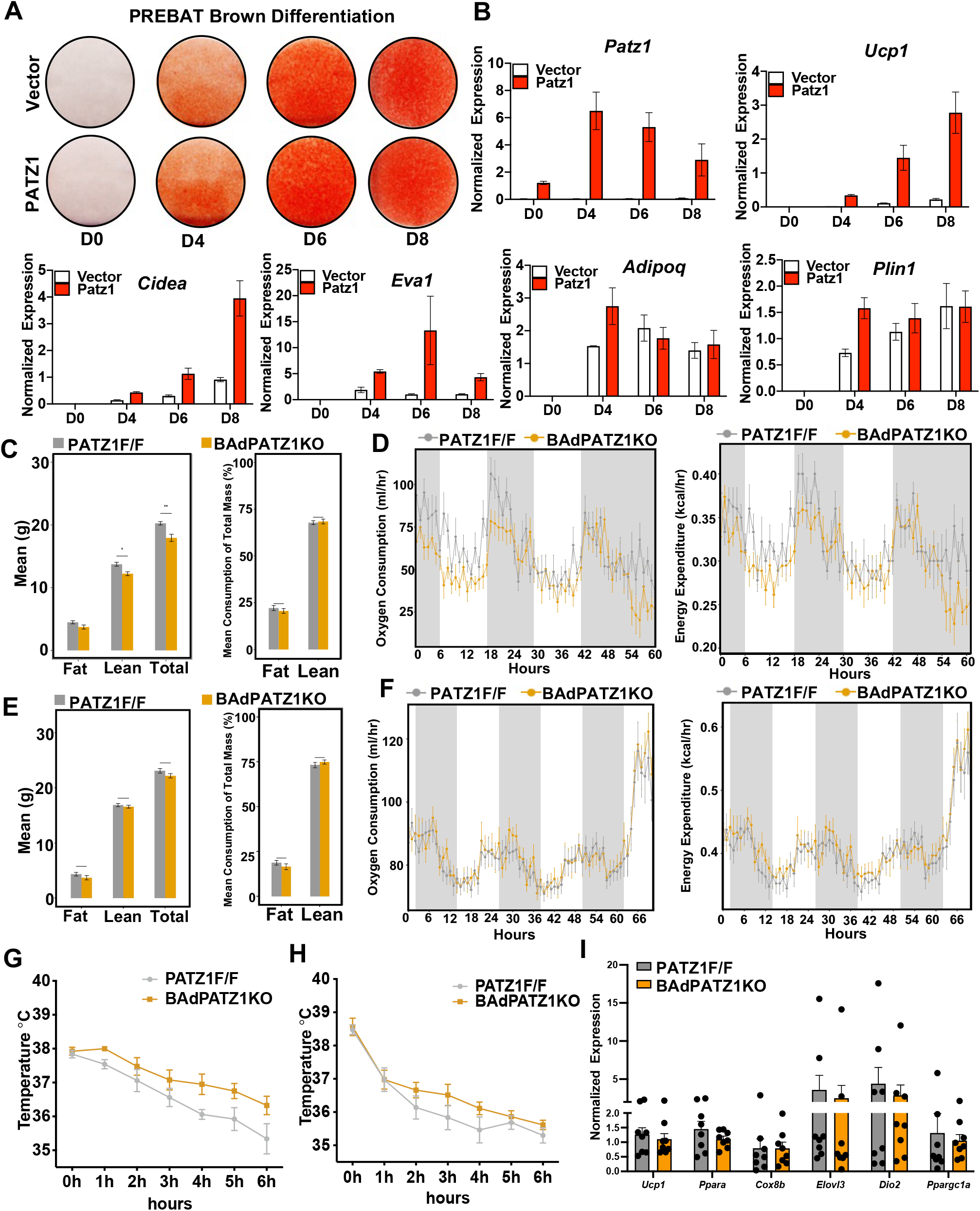
PATZ1 is dispensable for BAT adipogenesis and thermogenesis. **A)** Oil-Red-O staining of immortalized brown preadipocyte (PREBAT) cells stably expressing Vector or PATZ1, brown-differentiated for indicated days. **B)** Real-time qPCR of indicated genes from Vector or PATZ1 overexpressing PREBAT cells brown differentiated for indicated days. **C)** Body fat, lean mass, and total mass in PATZ1F/F and BAdPATZ1KO mice at 11 weeks of age determined by EchoMRI. **D)** Oxygen consumption (ml/hr) and Energy Expenditure (kcal/hr) were analyzed by Sable Promethion System metabolic chambers. Twelve-hour light/dark cycles, 60-hour total duration; each light/grey bar represents 12-hour duration. **E)** Body fat, lean mass, and total mass in PATZ1F/F and BAdPATZ1KO mice at 24 weeks of age determined by EchoMRI. **F)** Oxygen consumption (ml/hr) and Energy Expenditure (kcal/hr) of mice in (E) were analyzed by Sable Promethion System metabolic chambers. Twelve-hour light/dark cycles, 60-hour total duration; each light/grey bar represents 12-hour duration. 0-60 h duration was at 22°C and last 6 h was at 4°C. **G and H)** Rectal temperature of males (G) and females (H) PATZ1F/F and BAdPATZ1KO mice exposed to 4°C for 6 hours. **I)** Realtime qPCR of indicated genes from BAT derived from cold-exposed PATZ1F/F and BAdPATZ1KO mice. For all the figures panels, n = 6-8 per group, and representative cohort from at least three independent studies. * p<0.05, **p<0.01, *** p<0.001, n.s. not significant.

BAT plays an important role in non-shivering thermogenesis during cold exposure. To test if PATZ1 deletion in BAT affects cold tolerance, we exposed both male (**Fig. 5G)** and female (**Fig. 5H)** BAdPATZ1KO and control mice to cold stress (4°C) for 6 hours and measured their rectal temperature. As shown in **Fig. 5G and 5H**, both genotypes showed similar changes in temperature, and we did not notice heightened or attenuated cold tolerance in BAdPATZ1KO mice compared to controls. In agreement with our cold exposure and indirect calorimetry data, BAT thermogenic gene expression was comparable between BAdPATZIKO and controls (**Fig. 5I**). These data suggest that PATZ1 primarily affects the function of WAT.

### Global chromatin and transcriptomic dynamics influenced by PATZ1

To characterize the DNA binding dynamics of PATZ1 during adipocyte differentiation, we analyzed genome-wide binding of PATZ1 using ChIP-seq (See Methods) during D0 (D0; 2 days of insulin and GW1929) and D5 (D5; 2 days of insulin and GW1929, 2 days of adipogenic cocktail, and 1 day of insulin and GW1929) of 10T1/2 adipogenic differentiation. Motif enrichment analysis of the ChIP-seq data showed highest enrichment of the PATZ1 motif, validating our immunoprecipitation method (**Fig. 6A**). Further analysis showed that PATZ1 bound to a consensus site of a stretch of guanines (G) and that genome-wide PATZ1 binding was enriched at intergenic and intronic regions (**Fig. 6A**). The percentage of promoter occupancy of PATZ1 markedly decreased from 20% at D0 to 9% at D5, indicating a substantial loss of PATZ1 promoter interactions during differentiation (**Fig. 6A**). A UCSC genome browser view of top PATZ1 peaks showed that PATZ1 bounds to its own promoter region at D0, but this binding was nearly completely lost at D5. These data suggest that PATZ1 regulates its own levels during early differentiation (**Fig. 6B**).

**Figure 6.**
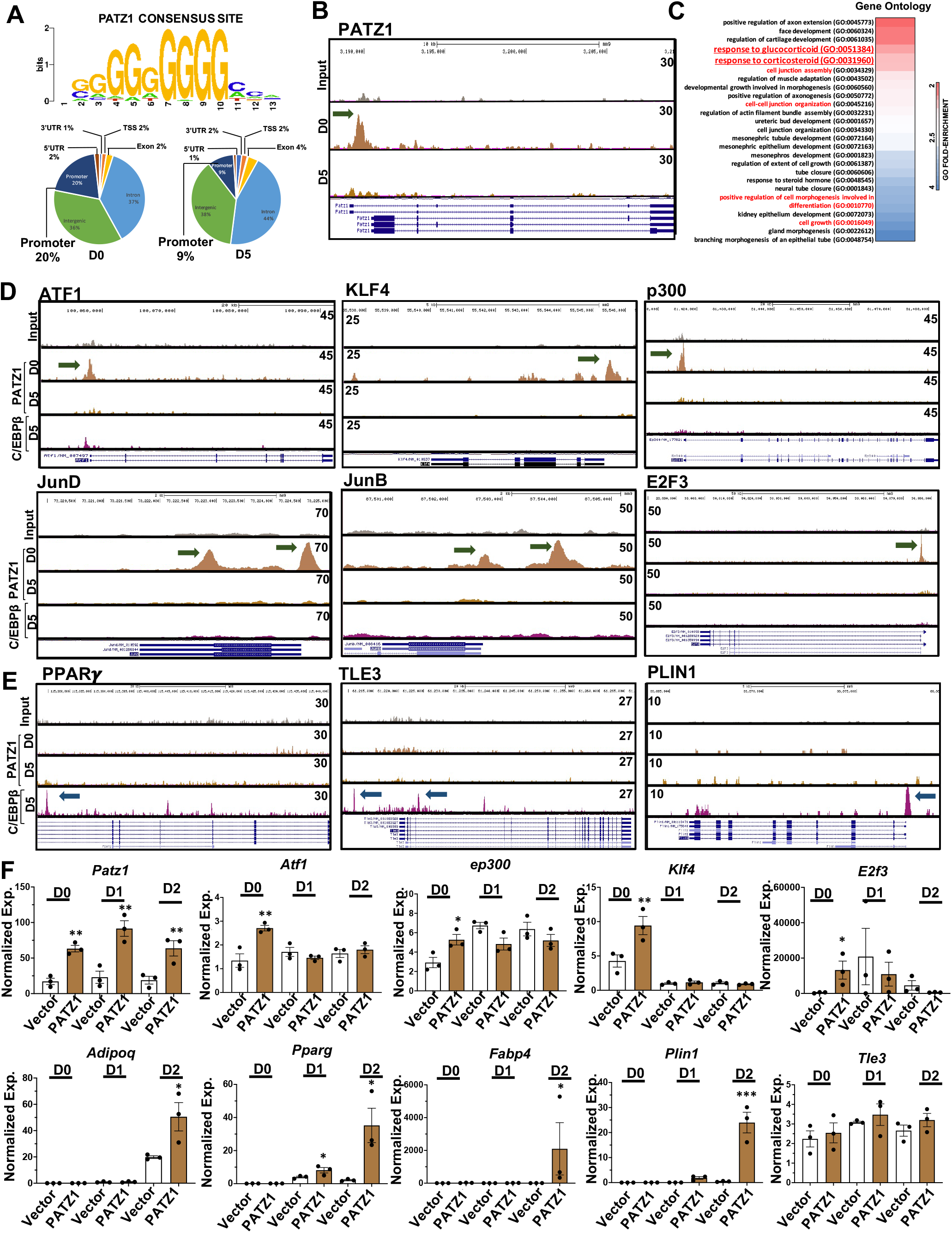
PATZ1 binds to the promoter regions during early adipocyte differentiation. **A)** ChIP-seq data derived consensus binding site of PATZ1 (top) and global occupancies of PATZ1 from PATZ1 ChIP-seq in 10T1/2 cells differentiated for D0 and D5 (bottom). **B)** BedGraph image peaks from PATZ1 ChIP-seq data showing control (input) and PATZ1 peaks at the promoter regions of PATZ1 locus on D0 and D5 differentiated 10T1/2 cells. **C)** Gene ontology on top PATZ1 peaks based on fold enrichment. **D)** BedGraph image peaks from PATZ1 and C/EBPß ChIP-seq data showing control (input) and PATZ1 and C/EBPß peaks at the promoter regions of indicated genes on D0 and D5 differentiated 10T1/2 cells. Numerical values in the BedGraphs are peak heights set in the UCSC genome browser. Green arrows show PATZ1 peaks. **F**) Real time qPCR of indicated genes from D0-D2 differentiated 10T1/2 cells expressing PATZ1 or vector. **, p<0.01; *, p<0.05.

Gene ontology on above-threshold PATZ1 peaks showed a glucocorticoid responsive promoter interaction, which is consistent with prior glucocorticoid receptor (GR) ChIP-seq studies showing shared enrichment of DNA interactions with PATZ1 ^26^ (**Fig. 6C**). Further analysis of top PATZ1 peaks showed that PATZ1 binds preferentially to genes known to play key roles in the early stages of adipocyte differentiation, including ATF-1 ^27^, AP-1 factors ^28^ and KLF4 ^6^, and E2F3 ^29^ (**Fig. 6D)**. We identified a robust peak on the promoter regions of p300/CBP ^30^, which are important for adipose formation (**Fig. 6D**). As a control, we included ChIP-seq for C/EBPß ^31^ to compare PATZ1 binding to a known transcriptional regulator at D5 of differentiation. Although PATZ1 caused an increase in the expression of markers of late differentiation such as *Pparg, Tle3 and Plin1*, we detected recruitment of C/EBPß, but not PATZ1 to the TSS of those genes (**Fig. 6E**) ^31^. Directed qPCR showed an increase in the expression of early adipogenic factors (*Atf1, ep300, Klf4, and E2f3*) in PATZ1 expressing cells, supporting our ChIP-seq data (**Fig. 6F**). By D2 of differentiation, PATZ1 expression also caused increase in the transcription of late adipocyte genes such as *Pparg, Adipoq*, and *Plin1*. These results show that PATZ1 may play a role in the early phases of adipocyte differentiation, when major chromatin remodeling events occur ^32^.

### GTF2I interaction attenuates PATZ1 adipogenic function

Our results show that both DNA-binding and protein-interaction domains are important for PATZ1’s adipogenic function. Additionally, our genome-wide DNA-binding studies show that PATZ1 preferentially binds to the promoter region of adipogenic factors to mediate adipocyte differentiation. To further assess the role of protein interaction of PATZ1, we used an unbiased global PATZ1 IP-MS to identify the PATZ1 interactome in differentiated 10T1/2 cells. Among all the interacting partners, GTF2I showed i) the highest protein abundance index (EmPAI), ii) the highest PATZ1/IgG abundance ratio, iii) the highest abundance p-value (**Fig. 7A**). Similar to PATZ1, GTF2I is expressed by APCs and adipocytes WAT ^20^ and is upregulated during adipogenesis, indicating a possible role for GTF2I in adipocyte differentiation (**Fig. 7B and 7C**). To test if GTF2I was regulated by PATZ1, we measured GTF2I levels in control and PATZ1 overexpressing differentiated 10T1/2 cells and found that GTF2I was consistently upregulated during differentiation (**Fig. 7D**). However, GTF2I levels were comparable between PATZ1 and vector cells (**Fig. 7D**). To test the role of GTF2I in PATZ1 adipogenic function, we knocked down GTF2I in PATZ1-overexpressing 10T1/2 cells and differentiated them into adipocytes until D2 and D4. As shown in **Fig. 7E**, we achieved an almost 100% depletion of GTF2I in *GTF2I* siRNA-treated cells compared to control- or PATZ1-expressing cells. Surprisingly, transcript analysis using qPCR from four independent experiments showed that PATZ1-mediated upregulation of adipocyte genes such as *Adipoq, Pparg, Cfd, Plin1, Fabp4* were markedly augmented by knocking down GTF2I (**Fig. 7F and 7G**). GTF2I can function as a transcriptional repressor or activator, therefore, our data suggest a model in which GTF2I could form repressive interactions with PATZ1 at the promoter regions of adipogenic regulators to dampen differentiation.

**Figure 7.**
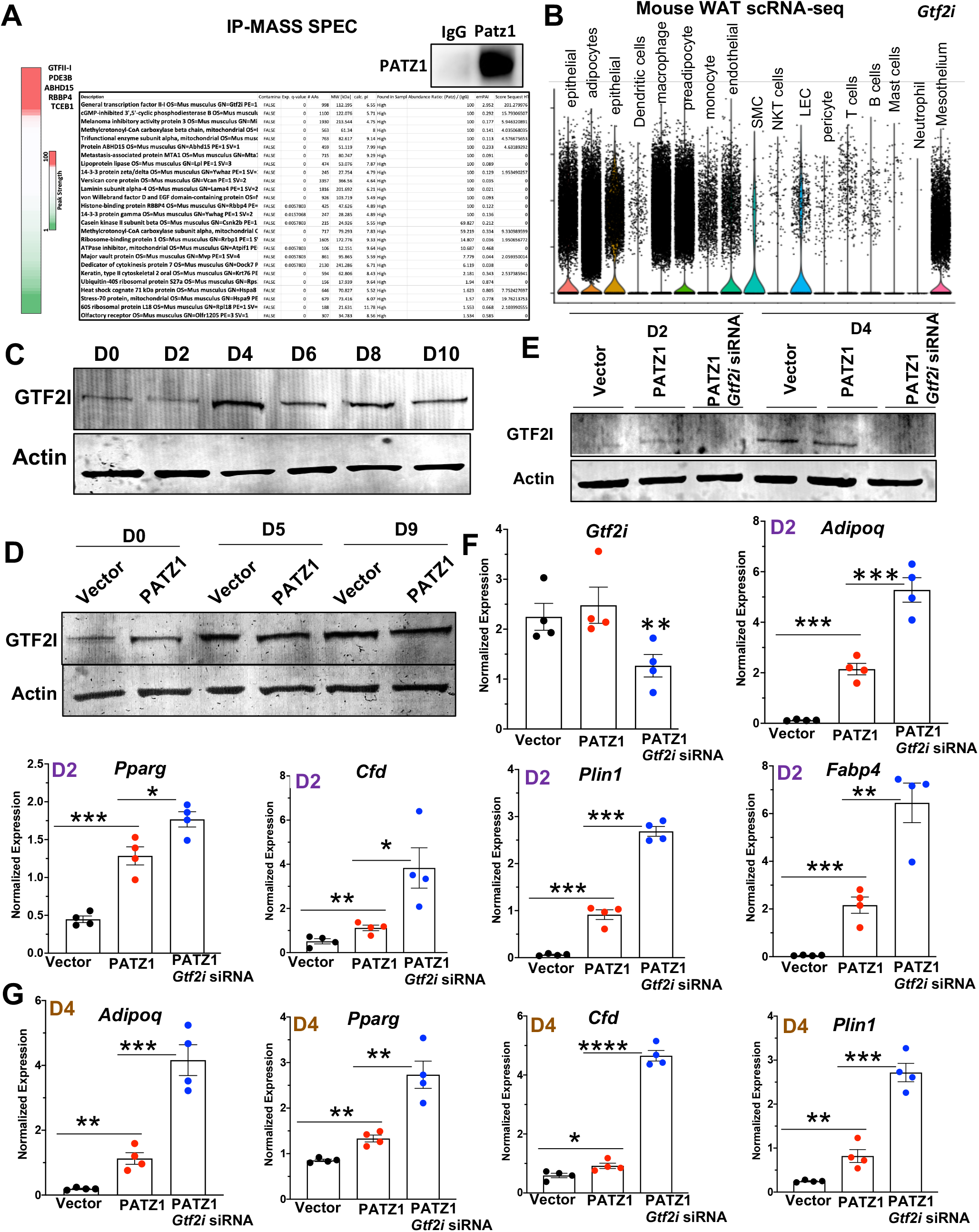
GTF2I attenuates PATZ1 adipogenic function. **A)** (Top) Immunoblot showing immunoprecipitation of endogenous PATZ1 from D2 differentiated 10T/2 cells. IgG was used as a control. (bottom) Heatmap and LC/MS mass spec data showing abundance of indicated protein peptides in PATZ1 pulldown experiment. **B)** Violin plot of scRNA-seq ^20^ cluster analysis showing levels of GTF2I mRNA in mouse WAT. **C)** Immunoblot analysis of GTF2I protein during differentiation of 10T1/2. Actin was used as a loading control. **D)** Immunoblot analysis of GTF2I protein during differentiation of 10T1/2 expressing Vector or PATZ1. Actin was used as a loading control. **E)** Immunoblot analysis of GTF2I protein during differentiation of 10T1/2 expressing Vector, PATZ1, or PATZ1 transduced with *GTF2I* siRNA. Actin was used as a loading control. **F and G)** Realtime qPCR of indicated gene from D2 (F) and D4 (G) differentiated 10T1/2 cells expressing Vector, PATZ1, or PATZ1 transduced with *GTF2I* sIRNA. Data are from four independent experiments. ****, p<0.0001; ***, p<0.001; **, p<0.01; *, p<0.05.

## Discussion

Adipogenesis is a highly orchestrated event involving interplay between transcriptional networks. In the present study, we have delineated a role of PATZ1 in transcriptional events controlling adipocyte differentiation. PATZ1 is expressed in both SVF and adipocyte portions of WAT. PATZ1 is not a PPARγ target gene, and its level remains relatively constant during adipocyte differentiation. Our data show that PATZ1 promotes adipogenesis through a mechanism dependent on DNA binding and protein interaction. Knockout of PATZ1 in adipose tissues protects mice from diet-induced obesity. Mechanistically, PATZ1 binds to the promoter regulatory regions of early adipogenic factors. We also identified another transcription factor affecting adipogenesis, promoter-binding transcription factor GTF2I, that attenuates PATZ1 adipogenic function.

Previous work has linked zinc finger proteins (ZFPs) to adipogenesis. Early commitment factors such as ZFP423 are known to drive *in vivo* adipogenesis and maintain white adiposity ^4,33,34^. ZFP467 suppresses osteogenesis and promotes adipogenesis of fibroblast-like progenitors by enhancing the expression of another master regulator of adipogenic transcription, CCAAT/enhancer binding protein-α (C/EBPα) ^35^. ZFP KLF5 acts as a co-activator of C/EBPβ, which then enhances PPARγ and C/EBPα transcription ^36^. Architecturally similar ZFPs containing BTB/POZ domains, such as BCL6 and PLZF, have also been implicated in adipogenesis ^37-39^. Our discovery of PATZ1 as a new adipogenic transcription factor adds to the diverse repertoire of ZFP transcription factors in regulating various stages of adipogenesis.

Genome-wide chromatin accessibility approaches, transcriptomics (RNA-seq and scRNA-seq), and chromatin immunoprecipitation-seq (ChIP-seq) have provided mechanistic insights into the role of TFs driving initial stage of adipogenesis ^40^. DNase I hypersensitive (DHS) regions based on ultradeep sequencing and ChIP-seq have revealed genomic foot printing and hotspots at enhancer and super-enhancers of PPARγ occupied by early adipogenic factors such as AP1, GR, ATFs and KLFs ^41^. This approach has revealed a network of the first wave of commitment factors that modify the chromatin environment of terminal factors like PPARγ to enable terminal adipocyte differentiation. Genome-wide glucocorticoid receptor (GR) Chip-seq in differentiated 10T1/2 cells ^26^ and FAIRE-seq (Formaldehyde-Assisted Isolation of Regulatory Elements-seq) in differentiated 3T3-L1 cells ^42^ showed abundance of the PATZ1 consensus region, hinting towards a plausible role of PATZ1 in adipose biology. We performed an unbiased ChIP-seq study to probe for PATZ1 DNA binding capacity during adipogenesis. We observed that the PATZ1 cistronic region includes GC-rich motifs. PATZ1 promoter occupancy revealed marked changes between D0 and D5. Loss of PATZ1 promoter binding at D5 of adipocyte differentiation despite constant PATZ1 expression levels suggests a differentiation-dependent DNA-binding capacity of PATZ1. PATZ1-binding peaks were primarily present at the promoter regions of genes known to play key roles in the early stages of adipocyte differentiation, including ATF-1 ^27^, AP-1 factors ^28^, KLF4 ^6^, and E2F3 ^29^. We also identified a robust peak of PATZ1 binding on the promoter region of p300, which is known to be critical for adipocyte formation ^30^. Further, our directed qPCR studies showed an increase in the expression of these factors in PATZ1 expressing cells, directly supporting the role of PATZ1 binding in increasing their gene expression. Of particular interest, PATZ1 increased the expression of markers of late differentiation (*Pparg, Tle3 and Plin1)*, but we did not detect recruitment of PATZ1 to the TSS of those genes (**Fig. 4A**). These findings implicate PATZ1 actions in the first wave of early regulators that ultimately lead to increase in terminal differentiation. Our results suggest that PATZ1 plays a key role in the early and commitment phases of adipocyte differentiation, when major chromatin remodeling events occur ^32^.

Our animal data also suggest a role of PATZ1 in late adipogenesis, since, *Adipoq-*Cre mouse model will not deplete PATZ1 in uncommitted APCs and WAT mesenchymal stem cells (MSCs). Adipose tissue deletion of the *Patz1* gene resulted in decreased WAT mass and protection from diet-induced obesity in both sexes fed a HFD (60% kcal fat). These data suggest that PATZ1 does not contribute to a sex-specific adipose phenotype and further demonstrates PATZ1 also affects late stage adipogenesis in vivo.

To better understand how PATZ1 contributes to adipogenesis, we performed PATZ1 IP-MS on differentiated 10T/2 cells. Among all the interacting partners, GTF2I showed the highest abundance in the PATZ1 complex. GTF2I is a transcriptional factor involved in both basal transcription and signal-induced transduction activation or repression ^43^, therefore, GTF2I is not considered a general transcription factor but rather a protein that regulates transcription of a selected number of genes ^44^. Similar to PATZ1, GTF2I can both inhibit and activate genes depending on stimuli and cellular context. Another member of the GTF2I family of TFs, GTF2IRD1, is known to bind PRDM16 and suppress the adipose tissue fibrosis gene program ^45^. ChIP-seq studies have shown a high preference of GTF2I to bind to the promoter regions of its target genes ^46^, and its occupancy overlaps with activating H3K4me3 peaks that mark transcriptionally active regions. Strikingly, genome sequences under GTF2I bound peaks are enriched for motifs for transcriptional regulators including PATZ1 ^46^. We showed that GTF2I is upregulated during adipocyte differentiation. Surprisingly, we found that GTF2I interaction with PATZ1 could form a repressive complex to inhibit adipogenesis. PATZ1 overexpression did not cause upregulation of GTF2I, suggesting a potential interaction-based dynamic of PATZ1 and GTF2I in regulating adipogenesis. Overall, we have found PATZ1 and GTF2I as two regulators of adipogenesis.

## Methods

### Reagents, Plasmids, and Viruse

Dexamethasone (D2915), 3-isobutyl-1-methylxanthine (IBMX, I7018), PPAR*γ* agonist GW1929 (G5668) and Oil-Red-O (O0625) were from Sigma-Aldrich. Insulin (12585-014) was from Thermo Scientific Fisher. cDNA encoding PATZ1, NFXL1, NCOA5 was cloned from the cDNA library into pBabe puro retroviral vector using Gateways Cloning System (Thermo). PATZ1 domain deletion mutants (ΔPOZ, ΔZnF, and POZ/AT) were cloned out of WT PATZ1 cDNA using site-specific and domain spanning primers and cloned in the pBabe retroviral vectors. pBABE-plamids were transfected into retrovirus packaging Pheonix E cells for 48hrs. Target cells (immortalized beige/brown preadipocytes) were plated at 50% confluency 24hrs post-transfection. 48 h after transfection media from transfected Phoenix-E cells were harvested and spun down for 5min at 5000 rpm to pellet cells and debris. Retrovirus containing supernatant was carefully removed and plated onto target cells with 1:1000 polybrene for overnight. Next day, media was replaced with regular growth media and cells were incubated for additional 24 h. 4.5 μg/ml puromycin selection was performed to select for cells stably expressing PATZ1. *Patz1* shRNA sequences were designed using BLOCK-iT RNAi designer tool (Invitrogen). Sense and antisense oligos were annealed and cloned into pENTR/U6 plasmid (Invitrogen). Using LR recombination (Invitrogen), shRNA constructs were subcloned into a Gateway-adapted pBabe-Puro plasmid. Only sense strands are shown here. Adenovirus was amplified, purified, and tittered by Viraquest. PATZ1 was cloned in to pDONR221 entry vector using the Gateway system. The sequences in the entry clones were then transferred by LR recombination into pAd/CMV/V5-DEST Gateway vector for viral particle production. All adenoviruses including Ad-GFP and Ad-CRE were amplified, purified, and titered by Viraquest. *Gtf2i* siRNA (ON-TARGETplus SMRTPool, Cat# L-049166-00-0005) was purchased from Horizon Discovery and Lipofectamine2000 (ThermoFischer) was used for transfection.

### Animals

C57BL/6 WT male and female mice (#000664), *Adipoq*-CRE (#028020), and *Myf5-* Cre (#007893) were acquired from Jackson Laboratory. *Patz1*^*f/f*^ (PATZ1 F/F) mice were generated as described previously^47^. All mice were maintained in a pathogen-free barrier-protected environment (12:12 h light/dark cycle, 22°C-24°C) at the Mount Sinai animal facilities. Animal experiments were conducted in accordance with the Mount Sinai Institutional Animal Care and Research Advisory Committee.

### Cell Culture

Murine preadipocyte cell lines 3T3-L1 and 10T1/2, and primary SVFs were cultured in DMEM supplemented with 10% fetal calf serum (FCS) for 3T3L1 and 10% fetal bovine serum (FBS) for 10T1/2. For *in vitro* differentiation, preadipocytes were grown to confluence in DMEM with 10% FBS plus insulin (5 μg/ml). Confluent cells were induced to differentiate with dexamethasone (1 μM), IBMX (0.5 μM), insulin (5 μg/ml), and with or without GW1929 for 2 days, followed by insulin and GW1929 alone. Growth media was exchanged every 2 days during differentiation. All stable cells were generated using the pBabe retroviral system ^48^. Retrovirus was obtained by transfection of Phoenix E cells with the pBabe vector and harvest of growth media 72 h later. Preadipocytes were transduced with retrovirus overnight and selected with antibiotics. For adenoviral overexpression of GFP or PATZ1, differentiated 10T1/2 cells expressing hCAR were transduced with Ad-GFP or Ad-PATZ1 on indicated days for 24 hrs and cells were harvested for transcript analysis by qPCR. For adenoviral CRE mediated knock down of PATZ1, SVFs from iWATs of PATZ1F/F mice were infected with Ad-GFP or Ad-CRE for 24h and differentiated into white adipocytes and samples were harvested on D0 or D7. Frozen primary human ADSCs were derived from abdominal scWAT as described previously^49^ and were traduced with Vector or PATZ1 retrovirus and differentiated using published protocol^49^. For glucose study, 10T1/2 cells were maintained in DMEM without glucose and supplemented with 2% FBS. Cells were then treated with various concentrations of glucose (Corning) and harvested 6h or 16h (ON) after glucose treatment. For GTF2I knockdown experiments, final concentration of 40 nM of *Gtf2i* siRNA was transfected in a well of a 12 well plate (Corning) consisting of confluent PATZ1 overexpressing 10T1/2 cells for 24h. 48h post-transfection cells were differentiated into white adipocytes for 2 days or 4 days.

#### Isolation of and immortalization of primary white and brown adipocytes

C57BL/6 WT male mice (8–10-week-old) were euthanized in isoflurane chamber. Mice were sprayed thoroughly with 70% EtOH. 100-300 mg of inguinal WAT (iWAT) or BAT were dissected and placed on sterile 6-well tissue culture plate with ice-cold 1XPBS. Fat pads were blotted on a napkin to removed excess liquid. Tissues were cut with scissors and minced using blade. Minced fat pads (600-800 mg) were placed in a15ml conical tube containing 3 ml of digestion buffer (PBS, 1.5 U/ml Collagenase D (iWAT) or collagenase B (BAT), 2.4 U/ml Dispase II, 10 mM CaCl_2_) and incubated at 37°C for 45 min with gentle shaking. Inside tissue culture hood, 10-15 ml of plating media DMEM/F12 with glutamax supplemented with 15%FBS and 1% pen/strep was added to digested solution and slowly resuspended 5 times. The digestion mixture was passed through 100 μm cell strainer and centrifuged at 500 x g for 10 mins at room temperature. Supernatant was carefully decanted and pellet resuspended in 10 ml of plating media and passed through 40 μm cell strainer. Filtered suspension was spun down again at 500 x g for 10 mins at room temperature. Supernatant was decanted and pellet resuspended in 6ml plating media and plated onto collagen-coated plates. After overnight incubation, media was changed every other day until the cells reached 70% confluency (∼3-4 days post-harvest). White and brown stromal vascular fractions (SVF) were with differentiated into beige/brown adipocytes using protocol mentioned above or immortalized. For immortalization, retrovirus-expressing largeT-antigen in pBABE-hygromycin vector was generated as mentioned above. Virus was added to target cells and selected with 600 μg/ml hygromycin to make immortalized cells.

### Gene Expression Analysis

Total RNA was isolated using TRIzol reagent (Invitrogen) and reverse transcribed with the High-Capacity cDNA synthesis kit (Thermo). cDNA was quantified by real-time PCR using SYBR Green Master Mix (Diagenode) on a Quant Studio 5 384 well plate. Gene expression levels were determined by using a standard curve. Each gene was normalized to the housekeeping gene 36B4 and was analyzed in duplicate. Primers used for real-time PCR are listed in Table S1.

### Protein Analysis

Whole cell lysate was extracted using RIPA lysis buffer (Boston Bioproducts) supplemented with complete protease inhibitor cocktail (Roche). Proteins were diluted in Nupage loading dye (Invitrogen), heated at 95°C for 5 min, and run on 4–12% NuPAGE Bis-Tris Gel (Invitrogen). Proteins were transferred to hybond ECL membrane (GE Healthcare) and blotted with PATZ1 (A10053, Abclonal), GTF2I (PA5-17642, Thermo), Hmg1 (ab18256, Abcam), actin (A2066, Sigma), αTubulin (CP06, Calbiochem) antibodies.

### RNA-Seq

Total RNA was prepared as described ^50^. Strand-specific libraries were generated from 500 ng total RNA using the TruSeq Stranded Total RNA Library Prep Kit (Illumina). cDNA libraries were single-end sequenced (50bp) on an Illumina HiSeq 2000 or 4000. Reads were aligned to the mouse genome (NCBI37/mm9) with TopHat v1.3.3 and allowed one alignment with up to two mismatches per read. mRNA RPKM values were calculated using Seqmonk’s mRNA quantitation pipeline. All FPKMs represent an average from three biological replicates for in-vitro studies for library construction. A gene was included in the analysis if it met all of the following criteria: The maximum FPKM reached 10 at any time point, the gene length was > 200bp as determined by the DESeq2 package in R Bioconductor. P values were adjusted using the Benjamini-Hochberg procedure of multiple hypothesis testing^51^.

### Co-immunoprecipitation (Co-IP) and Mass Spectrometry

Cells expressing PATZ1 were lysed with ice-cold lysis buffer containing 50 mM Tris-HCl pH 7.5, 150 mM NaCl, 0.6% Triton X-100, plus complete protease inhibitor (Roche). Cell lysates were pre-cleared with empty Protein A/G PLUS agarose beads (Santa Cruz) and then immunoprecipitated with beads prelinked with anti-Flag M2 antibody (F1804, Sigma) overnight at 4°C. Beads were washed with 50 mM Tris-HCl pH 7.5, 150 mM NaCl, and 0.1% Triton X-100, then eluted with 3X Flag peptides (F4799, Sigma). In specific experiments, RNase cocktail enzyme (AM2286, Ambion) was used to treat cell lysates before immunoprecipitation. IP Eluents from Flag-PATZ1 cells and control cells were processed by UCLA Pasarow Mass Spectrometry Laboratory. The results were analyzed using the Mascot software.

### Mature Adipocyte and SVF Fractionation

Cells were washed with 1× PBS and incubated with TEN-buffer (10 mM Tris-HCl pH 7.5, 100 mM NaCl, 1 mM EDTA pH 8.0). Cells were allowed to swell on ice for 15 min in 10 mM HEPES, 10mM KCl, 0.1 mM EGTA, 0.1 mM EDTA, 1 mM DTT, plus complete protease inhibitor (Roche). Cells were then mixed with NP-40 alternative (Calbiochem) to a final concentration of 0.4%, incubated for 5 min, and centrifuged at 12,000 × *g* for 5 min. Supernatant was collected as the cytoplasmic fraction. Nuclear pellet was resuspended in 20 mM HEPES, 420 mM NaCl, 1.5 mM MgCl_2_, 0.2 mM EDTA, 25% glycerol, 1 mM DTT, plus complete protease inhibitor. After a 15-min incubation, nuclear extracts were spun at 12,000 × *g* for 5 min and the supernatant was saved for future analysis. For RNA analysis, all the buffers above were supplemented with 10 mM ribonucleoside-vanadyl complex (NEB) to inhibit ribonucleases. RNA was extracted from the nuclear and cytoplasmic fraction using Purelink RNA mini kit (Ambion).

#### Chromatin immunoprecipitation (ChIP)-seq (ChIP-Seq)

ChIP experiments were performed according to standard protocols ^52,53^. Lysed cells were sonicated using a Bioruptor (Diagenode) according to the manufacturer’s protocol, and chromatin was immunoprecipitated with antibodies against PATZ1 (sc-390577, Santa Cruz Biotechnology) and IgG (PP64, Millipore) overnight at 4°C in the presence of Protein A beads (GE Healthcare). ChIP-Seq libraries were prepared using the Kapa LTP Library Preparation Kit (Kapa Biosystems). ChIP-Seq was performed as described ^31,50^. Reads were aligned to the mouse genome (NCBI37/mm9) with Bowtie2. Uniquely mapped reads were used for peak calling and annotation using HOMER ^54^. Peaks were called if they passed a false discovery rate of 0.01 and were enriched over input. Peaks were annotated to the nearest TSS.

### Cold exposure studies

For 4°C cold exposure experiment, PATZ1 F/F or BAdPATZ1KO mice at 8-10 weeks of age were singly or doubly housed at 4°C in a non-bedded cage with access to food and water for the time points indicated in the figure legend. At the end of the experiment, BATs were resected for gene expression analysis. Core body temperature of WT and *Il10*^-/-^ mice was measured at room temperature using rectal probe (BAT-10) purchased from Physitemp.

### High fat diet studies

For diet study, 10 weeks of age AdPATZ1KO and PATZ1F/F mice were fed a 60% high-fat diet (Research Diets) for the indicated times. Mice weights and body composition were measured every week and food was replaced weekly.

### Indirect Calorimetry and Body Composition Measurements

Indirect calorimetry was performed using a Promethion Metabolic Chambers (Sable Systems). Animals were placed individually in chambers for 3 consecutive days at ambient temperature (22.5°C) or 6h in cold temperature (4°C) with 12 hr light/dark cycles. Animals had free access to food and water. Respiratory measurements were made in 6 min intervals after initial 7-9 hr acclimation period. Energy expenditure (EE) was calculated from oxygen consumption (VO2) and respiratory exchange ratio (RER) using the Lusk equation, EE in Kcal/hr = (3.815 + 1.232 X RER) X VO2 in ml/min. Food intake was measured in metabolic chambers. Body composition (fat and lean mass) was determined using EchoMRI Body Composition Analyzer. Indirect calorimetry data were analyzed by CALR web-based software^55^.

### Tissue hematoxylin and eosin (H&E) staining and immunohistochemistry

Tissues (4-5 microns thickness) were placed in cassettes and submerged in 10% formalin solution overnight. Tissue cassettes were washed with tap water for 15 minutes and stored in 70% EtOH at room temperature. Paraffin embedment and H&E staining was performed at the Pathology Core Laboratory at Mount Sinai.

### Mouse and human population-based investigation of *Patz1*

Hybrid mouse diversity panel data was analyzed from 106 inbred strains fed a High-fat high-sucrose diet for 8 weeks as previously described ^56,57^. Mouse adipose tissue global expression was analyzed using Affymetrix HT_MG430A arrays and overlaid with phenotypic measurements. Human adipose tissue expression arrays using Affymetrix U219 microarray and phenotypic data was analyzed from the Metabolic Syndrome in Men (METSIM) study ^24,25^. All correlations were assessed from the midweight bicorrelation coefficient and corrected p-value using the R package WGCNA ^58^.

## Acknowledgements

P.R is supported by R00DK114571, NIDDK-supported Einstein-Sinai Diabetes Research Center (DRC) Pilot & Feasibility Award, and Diabetes Research and Education Foundation (DREF) Grant # 501 (PR). M.S. is supported by NIH-NIDDK grants DK130640 and DK097771. W.E. has been supported for work on MAZR/PATZ1 by the Austrian Science Fund projects P19930 and P23641. P.T. is supported by DK120851. The funders had no role in study design, data collection and interpretation, or the decision to submit the work for publication.

## Contributions

S.P. and P.R. performed animal and in vitro differentiation experiments under the supervision of P.T. and P.R. PATZ1 F/F mice were generated by S.S and W.E. P.R. did initial in vivo and in vitro studies under the supervision of P.T. N.S. and S.S. performed immunoblot and qPCR under the supervision of P.R. P.R. and J.W. performed ChIP-Seq and IP-MS under the supervision of J.P.W., P.T. M.S. supervised and processed the RNA-seq data. H.W and C.J.V designed and supervised the cDNA library screen under the supervision of P.T. P.T. and P.R. conceived the project and P.T. and P.R. wrote the manuscript with help from J.W., S.K.F., H.W., C.J.V., M.S., and W.E.

